# 3D reconstruction of odontoblast processes of the mouse molar and incisor

**DOI:** 10.1101/2020.06.21.163626

**Authors:** Ninna Shuhaibar, Arthur R. Hand, Mark Terasaki

## Abstract

Odontoblast processes are thin cytoplasmic projections that extend from the cell body at the periphery of the pulp toward the dentin-enamel junction. The odontoblast processes function in the secretion and assembly of dentin during development, participate in mechanosensation, and aid in dentin repair in mature teeth. Because they are small and densely arranged, their three-dimensional organization is not well documented. To gain further insight into how odontoblast processes contribute to odontogenesis, we used serial section electron microscopy to examine these processes in the predentin region of mouse molars and incisors. In molars, the odontoblast processes are tubular with a diameter of ~1.8 μm. The odontoblast processes near the incisor tip are similarly shaped, but those midway between the tip and apex are shaped like plates. The plates are radially aligned and longitudinally oriented with respect to the growth axis of the incisor. The thickness of the plates is approximately the same as the diameter of molar odontoblast processes. The plates have an irregular edge; the average ratio of width (midway in the predentin) to thickness is 2.3 on the labial side and 3.6 on the lingual side. The plate geometry seems likely to be related to the continuous growth of the incisor and may provide a clue as to the mechanisms by which the odontoblast processes are involved in tooth development.

## Introduction

The tooth is composed primarily of dentin, a calcified extracellular matrix consisting of approximately 90% collagen type I, with the remaining 10% consisting of several proteins, glycoproteins and proteoglycans, some of which are tissue-specific whereas many are also found in bone and other connective tissue matrices (Goldberg et al. 2011). Dentin supports the enamel, the non-living, hard surface layer of the crowns of the teeth, and also forms the roots of the teeth which anchor them via the periodontal ligament to the alveolar processes of the mandible and maxilla. In the central part of the tooth, surrounded by the dentin, is the pulp, which consists of a loose connective tissue containing blood vessels and nerves, as well as immune cells and a progenitor cell population (Farges et al. 2015; Harichane et al. 2011).

Odontoblasts make dentin. Their columnar-shaped cell bodies are located in a densely packed single layer at the pulp / dentin boundary. Lying between the odontoblasts and the mineralized dentin is a 15-20 micron thick region called predentin because it consists of newly-deposited extracellular matrix that is not yet calcified. As new dentin is laid down and the dentin layer increases in thickness, the odontoblasts remain at the pulp / dentin boundary so that the pulp gradually becomes reduced in volume during development of the tooth.

Each odontoblast has a projection from its distal (secretory or apical) end called the odontoblast process that is embedded in the dentin, and extends from the cell body to the dentin-enamel junction (DEJ). As the dentin increases in thickness, the odontoblast process elongates, although in fully formed teeth the process may retract from the DEJ (Pashley, 1996). The processes are thought to be essential for the initial deposition of the dentin matrix as well as calcification (Weinstock and Leblond 1973, 1974; Nagai and Frank 1974; Rabie and Veis 1995). Each odontoblast process is located within its own narrow channel within the dentin, called a dentinal tubule. The odontoblast processes are present throughout life and there is evidence that, along with the cell body and nerves, they form a mechanosensory system, detect damage due to dental caries and attrition, and are involved in dentin repair (see reviews by Pashley 1996; Sasaki and Garant 1996; Smith and Lesot 2001).

Due to their small dimensions and dense distribution, the three-dimensional organization of dentinal tubules / odontoblast processes has not been thoroughly documented. We used the automated tape collecting ultramicrotome (ATUM) method for collecting serial sections for electron microscopy (Kasthuri et al. 2015) to characterize odontoblast processes in the predentin. These were examined in fully developed mouse molars and in the incisor.

## Results

Each side of the adult mouse mandible has three molars and one incisor. The molars develop during early life to a final shape, erupt postnatally, and are retained throughout life, whereas the incisors grow continuously. Transient stem cell niches (“enamel knots”) shape the molar (Jernvall and Thesleff 2000; Catón and Tucker 2009). A permanent stem cell niche at the incisor apex is the site of cell division (Kuang-Hsien Hu et al. 2014). Growth in this region pushes the older parts forward and the tip of the incisor is worn away during gnawing. Different positions along the length of the incisor are therefore at different stages of tooth development.

Two-month old mice were fixed by cardiac perfusion and the mandibles were removed. After decalcification, samples for electron microscopy processing were taken from the molar and from the tip and mid-incisor regions of the incisor (Figure 1). Serial sections (100-200 sections, 50-75 nm thick) were collected on an ATUM (automated tape collecting ultramicrotome; Kasthuri et al. 2015). The sections were imaged sequentially by scanning electron microscopy at a resolution that allowed us to discern cell boundaries and intracellular components. The odontoblast processes were clearly discernible in the uncalcified, predentin region between the mineralized dentin and the odontoblast layer at the periphery of the pulp (Figure 2).

**Figure 1.**
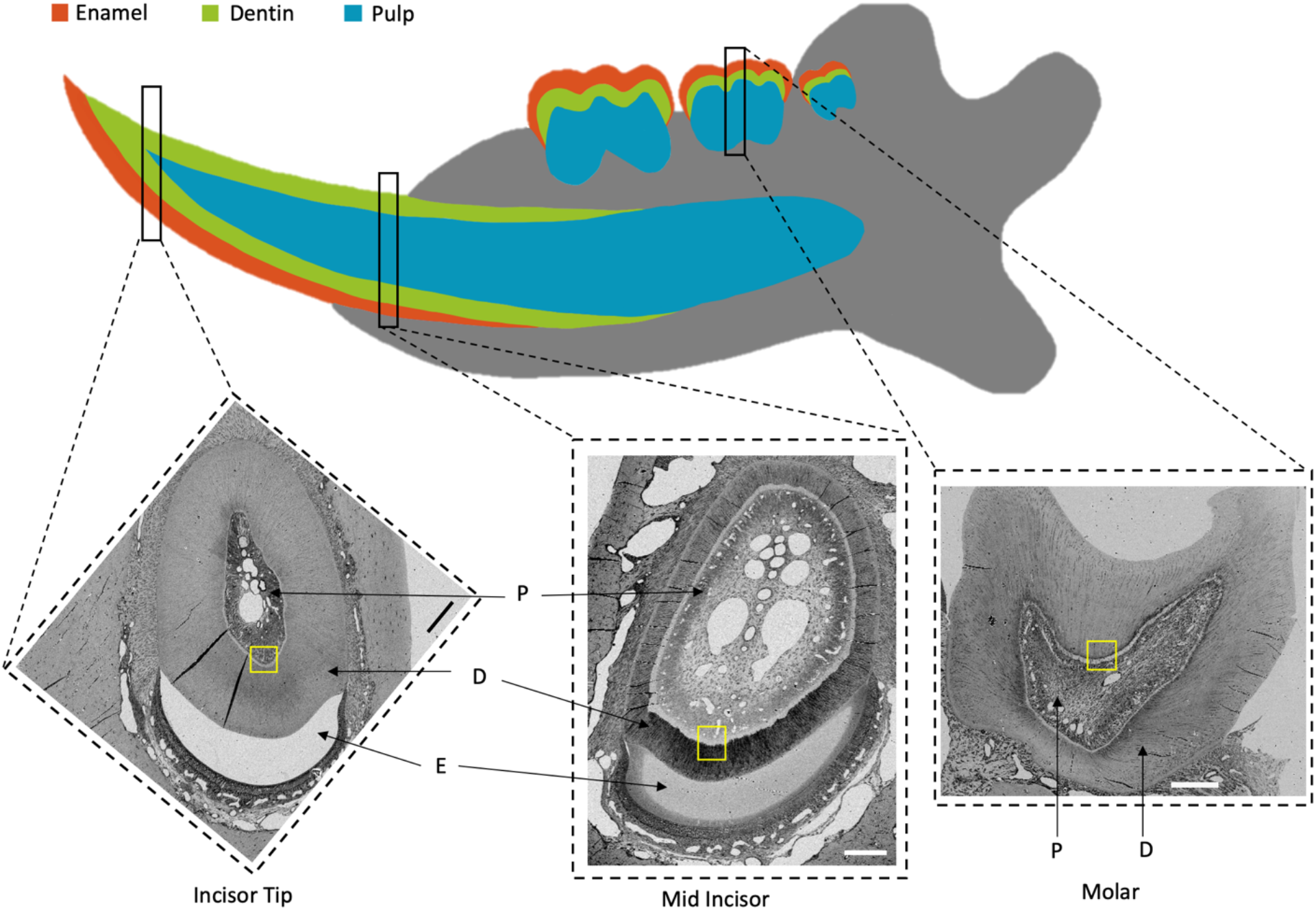
Diagram of the mouse mandible, showing the pulp (P), dentin (D) and enamel (E) regions of the molars and incisor. The insets are low magnification electron micrographs of the incisor tip, mid-incisor, and molar. The enamel of the molar was located in the top part of the inset but was dissolved away by the decalcification. The serial sections shown in Figure 2 were collected from the regions indicated by the yellow rectangles. Scale bars, for insets = 100 μm.

**Figure 2.**
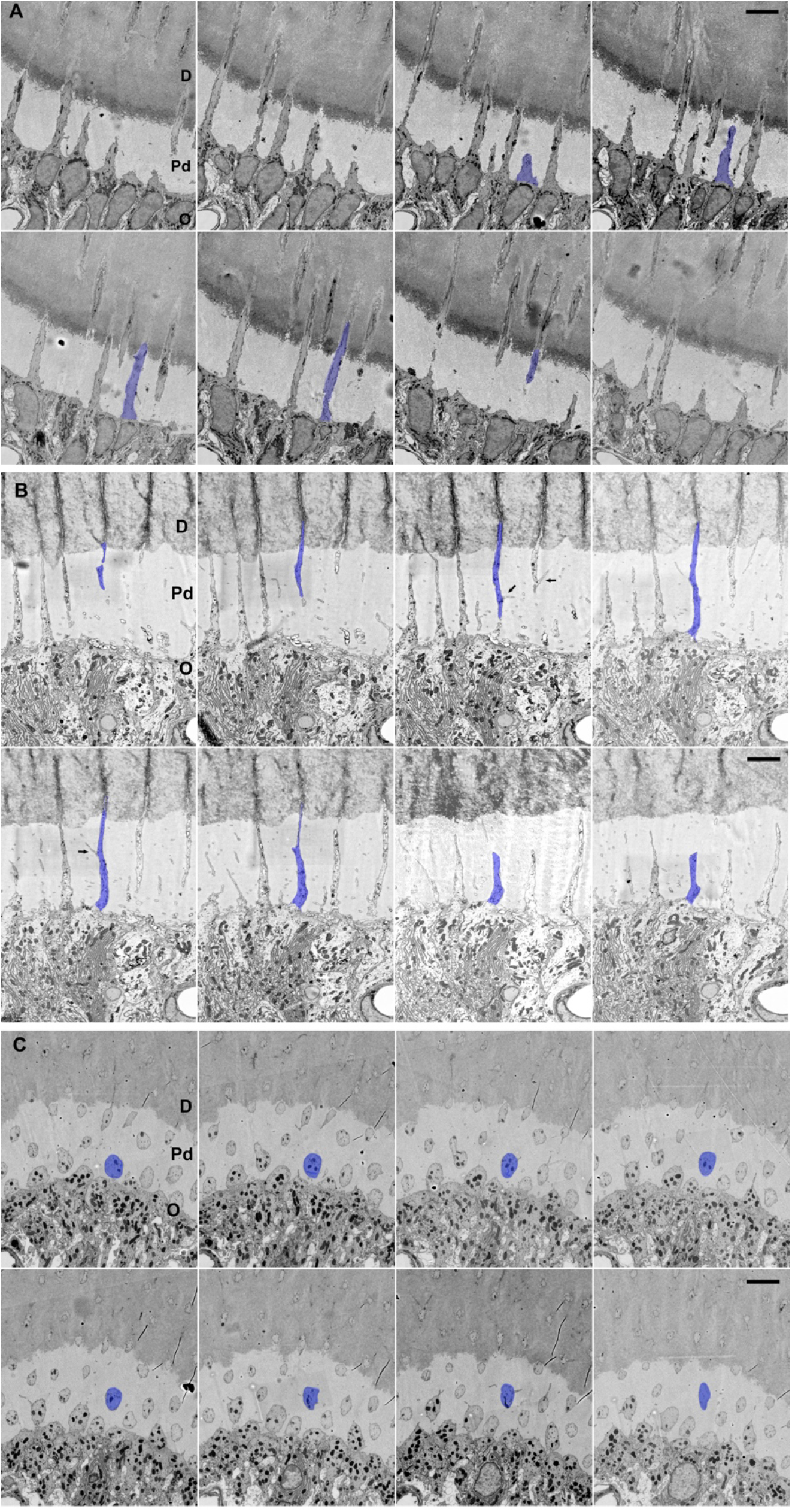
Serial section montages of A) molar, B) mid-incisor region, labial side, and C) incisor tip, labial side. The montages are printed at the same magnification. The individual sections from the molar and incisor tip were 50 nm thick and those from the incisor apex were 75 nm thick. The sections shown in each panel are ~570 nm apart. The total thickness of each series shown is ~4 μm. D, dentin; Pd, predentin; O, odontoblasts. The sections through the incisor tip processes in C) indicate that they have a tubular shape, but they appear to be much thicker than those in A) and B). This is explained by differences in sectioning planes and angles. The odontoblast cell bodies and predentin-dentin boundary are in parallel planes; the molar and mid-incisor sections (A, B) were cut perpendicular to these planes whereas the incisor tip sections (C) were cut obliquely to them. Note in A) and B) that the processes are thicker at their origin from the cell bodies and taper slightly as they approach the dentin. The sections in C) were cut closer to the pulp than to the predentin-dentin boundary. This explains why the cross sections appear much larger than those of A and B, while the reconstructions in Figure 3 and measurements in Table 1 show them to have a similar thickness. Scale bars = 4 μm.

From serial section data, the three-dimensional shapes and dimensions are best determined by segmentation and reconstruction. However, some salient features can be deduced by visual inspection of the images. A single odontoblast process is colored blue in each panel. The blue process is present in only 4 images of the molar (Figure 2A) whereas it is present in all 8 images of the mid-incisor (Figure 2B), indicating that the process is more elongated in the direction of sectioning. The sections through the incisor tip processes in Figure 2C indicate that they have a tubular shape. Three incisors from three different mice were examined and had the same elongated shape compared to the one molar that we examined which had a tubular shape.

The odontoblast processes were digitally segmented and reconstructed. In the molar and incisor tip, the odontoblast processes were cylindrically shaped (Figure 3A, C) and are slightly wider near the cell body than at the predentin-dentin boundary. In the mid-incisor region, the odontoblast processes have a plate-like shape (Figure 3B). The edges of the plates are irregular and generally tapered closer to the dentin. The plates are radially oriented with respect to the longitudinal axis of the incisor and aligned in the direction of overall growth (Figure 4).

**Figure 3.**
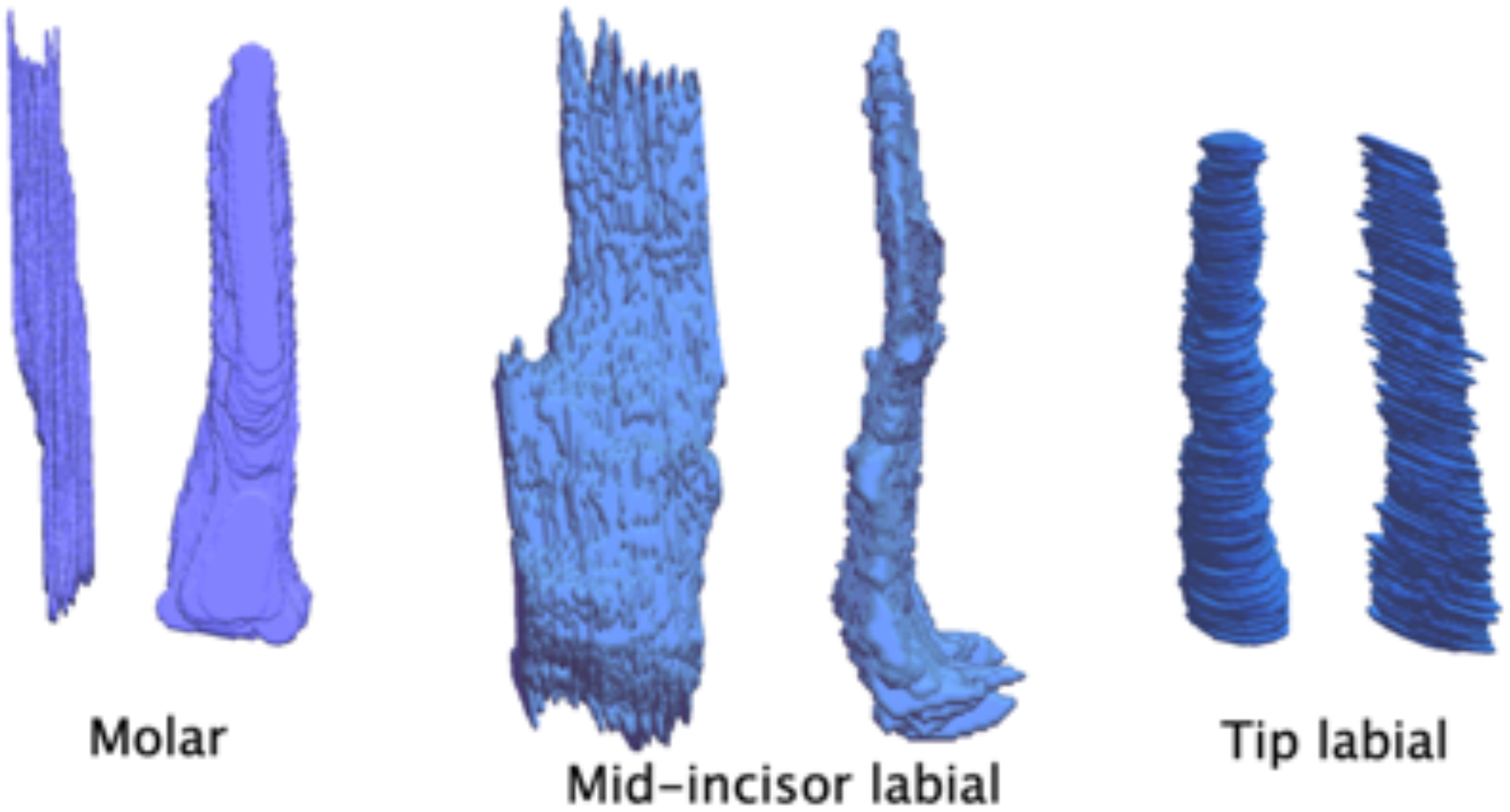
Reconstruction of an odontoblast process from Left) molar, Center) mid-incisor region, labial side, and Right) incisor tip, labial side. Each are viewed from two orthogonal directions. The cell bodies were located at the bottom of the figure, note that the processes taper as they approach the pre-dentin / dentin boundary at the top.

**Figure 4.**
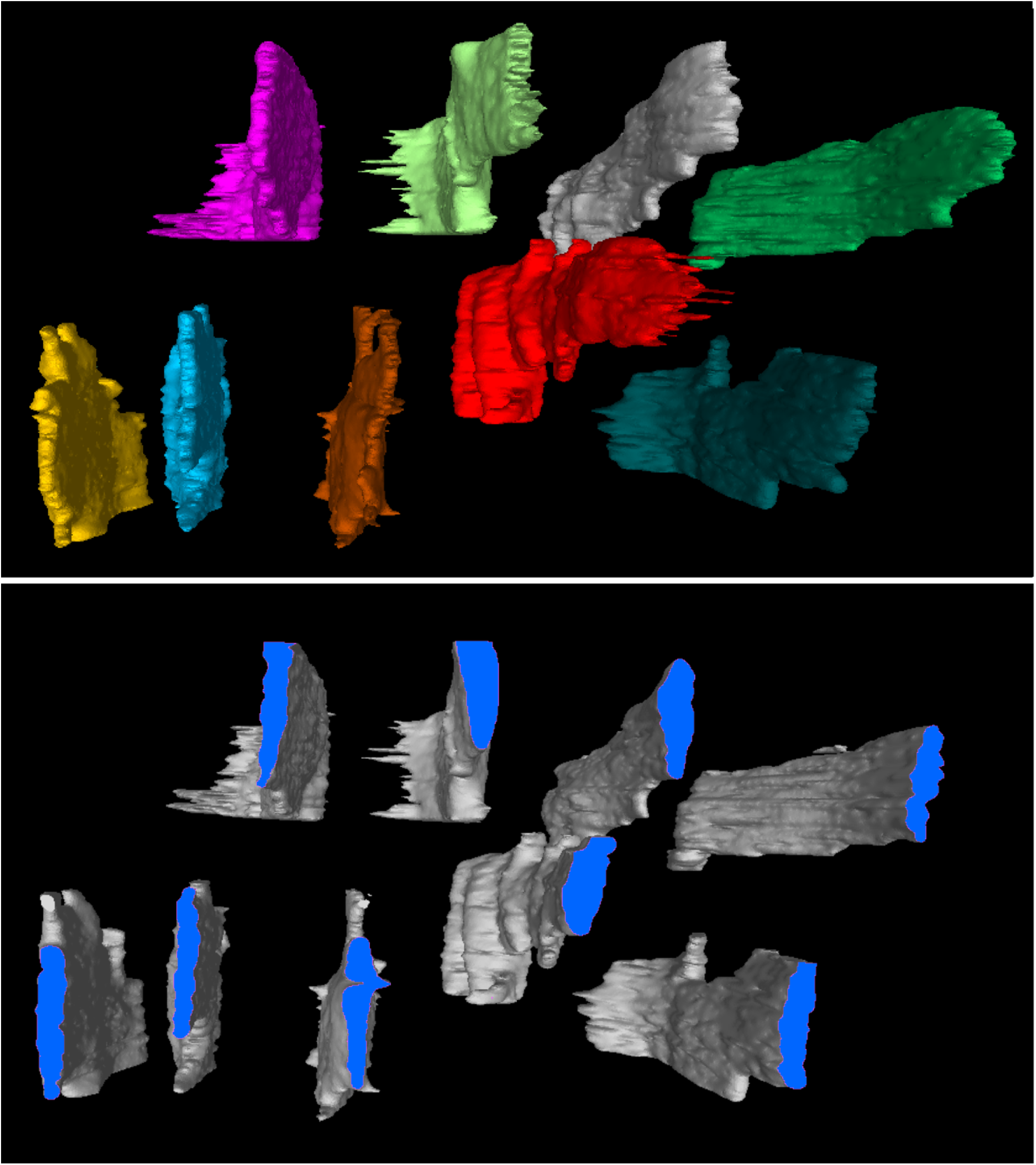
Reconstruction of a field of 9 odontoblast processes from the mid-incisor region, labial side. The cell bodies are located away from the viewer, while the predentin – dentin boundary is towards the viewer. The longitudinal axis of the incisor is vertical in the figure. The top panel shows the entire processes. The bottom panel shows them cut in cross-section in a plane midway between the odontoblasts and predentin – dentin boundary. The plates are all oriented in the same direction. Furthermore, the long axis of the plates is oriented parallel to the longitudinal axis of the incisor.

A virtual reality system was used to make measurements (see Methods). Cross-sections of the processes were measured midway in the predentin, and the largest (width) and smallest (thickness) dimension of the cross section were determined. The ratio of width to thickness indicates the shape of the cross-section. A ratio of one indicates a cylindrical shape and a ratio significantly greater than one indicates a platelike shape. The shape ratio for the processes of the molar was 1.14 ± 0.1. The shape ratio for the processes on all sides of the incisor tip (i.e., labial, lateral and lingual) were slightly higher (1.29 – 1.53), while the processes on all sides of the mid-incisor ranged from 2.26 to 3.62 (Table 1), with the lingual side being somewhat greater than the labial side.

**Table 1.**
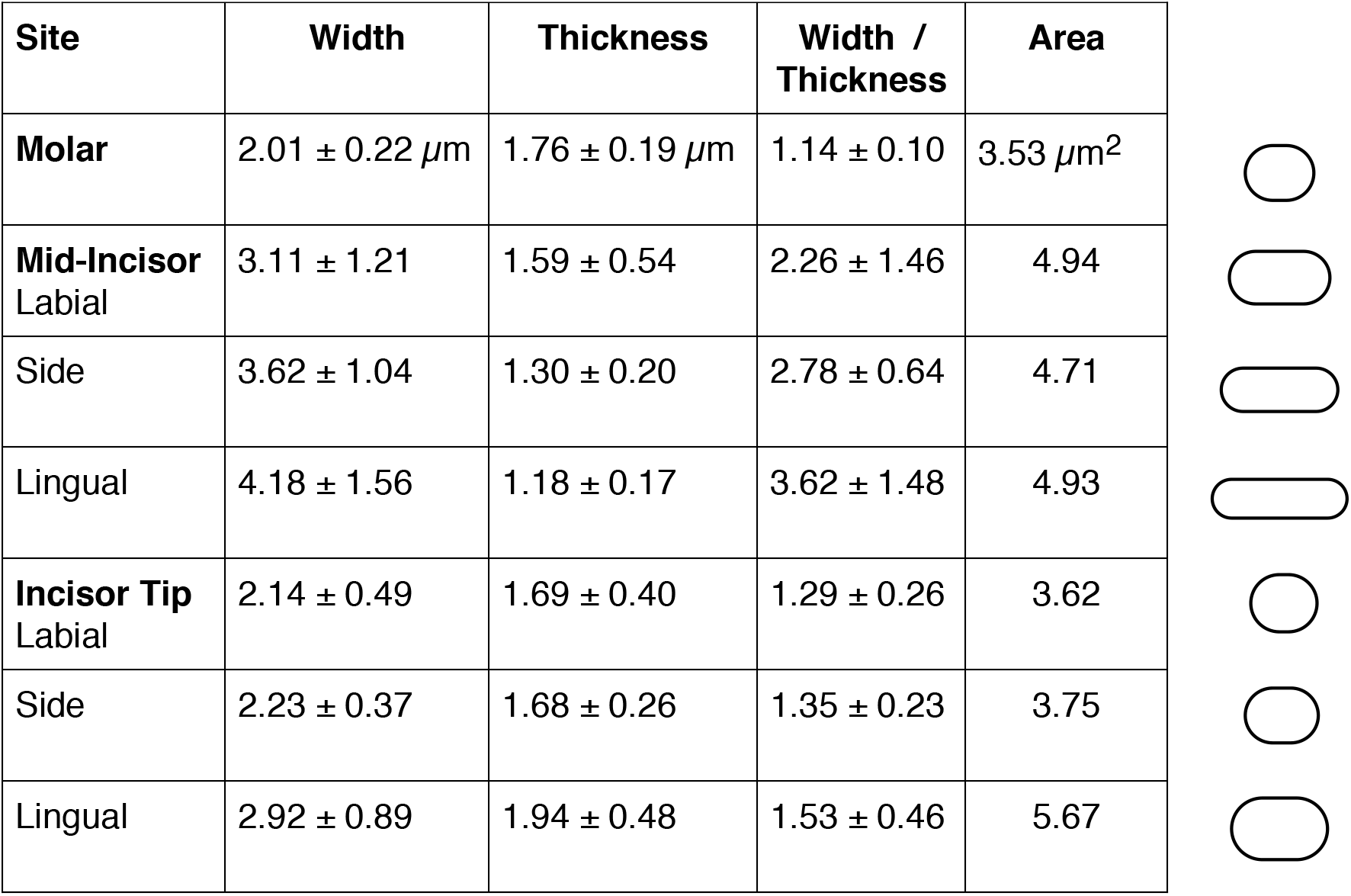
For each site, 12 odontoblast processes were measured. The measurements were made in the pre-dentin half way between the odontoblast layer and the predentin – dentin boundary. The ratio of width to thickness was measured individually in the 12 odontoblast processes then averaged. Cross sectional area is the product of average width and thickness. The diagrams on the right side of the table show visually the average width / thickness and illustrate how the odontoblast processes in the mid-incisor region resemble plates, while they resemble tubes in the molar and incisor tip.

| Site | Width | Thickness | Width / Thickness | Area |  |
| --- | --- | --- | --- | --- | --- |
| **Molar** | $2.01 \pm 0.22 \mu\text{m}$ | $1.76 \pm 0.19 \mu\text{m}$ | $1.14 \pm 0.10$ | $3.53 \mu\text{m}^2$ | |
| **Mid-Incisor**<br>Labial | $3.11 \pm 1.21$ | $1.59 \pm 0.54$ | $2.26 \pm 1.46$ | 4.94 | |
| Side | $3.62 \pm 1.04$ | $1.30 \pm 0.20$ | $2.78 \pm 0.64$ | 4.71 | |
| Lingual | $4.18 \pm 1.56$ | $1.18 \pm 0.17$ | $3.62 \pm 1.48$ | 4.93 | |
| **Incisor Tip**<br>Labial | $2.14 \pm 0.49$ | $1.69 \pm 0.40$ | $1.29 \pm 0.26$ | 3.62 | |
| Side | $2.23 \pm 0.37$ | $1.68 \pm 0.26$ | $1.35 \pm 0.23$ | 3.75 | |
| Lingual | $2.92 \pm 0.89$ | $1.94 \pm 0.48$ | $1.53 \pm 0.46$ | 5.67 | |

The average cross-sectional areas were calculated by multiplying average width by average thickness (Table 1). The areas fell into two groups, largely corresponding to shape and region. The more cylindrical processes of the molar and incisor tip had a smaller area, whereas plate-like processes of the mid-incisor region had larger areas. The incisor tip processes on the lingual side, which are less cylindrical, had a larger area.

As described previously (Sasaki and Garant 1996; Shahidi et al. 2015; Linde and Goldberg 1993), in both the molar and incisor, smaller processes branched from the sides of the main process (Figure 2B, arrows). These small processes often approached and appeared to contact similar processes from adjacent main processes.

## Discussion

Using recently improved methods for serial section electron microscopy, we documented the shapes of odontoblast processes in the predentin region of mouse teeth. In most textbook diagrams, the odontoblast processes are drawn having a tubular shape. We confirm this shape in molars and the tip of the incisor. Unexpectedly, the processes are plate-like in regions closer to the apex of the incisor. The short dimension (thickness) of the plates is comparable to the diameter of the tubule-shaped processes near the tip of the incisor. The plates are all radially oriented with respect to the longitudinal axis of the incisor, as if tubular processes had become extended sideways in the direction of incisor growth. The plates were somewhat wider on the lingual side of the incisor than labial side.

The plate-like shape of the incisor odontoblast process is most prominent for odontoblasts in the mid-incisor region, where matrix synthesis and secretion are occurring at a rapid rate (Takuma and Nagai 1971; Ohshima and Yoshida 1992). This shape would provide a greater surface area for secretion than a cylindrical shape. Near the incisor tip, where matrix synthesis and secretion rates are lower, the process is more cylindrical in shape. In the erupted molars examined in this study, secondary dentin matrix synthesis and secretion is occurring at very low rates, and the processes are nearly cylindrical. It would be interesting to examine the odontoblasts of developing molars, where active matrix formation is ongoing, to see if the processes have a platelike shape.

Another consideration is related to the different growth patterns of the molar and incisor. As dentin is laid down, the space occupied by the pulp gradually becomes reduced. In molars, the pulp is reduced in three dimensions. However, in the incisor, the pulp is reduced only in two dimensions. To a first approximation, the volume of dentin produced by a molar odontoblast is an elongated cone, while that by an incisor odontoblast is wedge-shaped, like a piece of pie.

The present results add significant details to the structure of the odontoblast process. Another report (Khatibi Shahidi et al. 2015), at the light microscopic level, described “odontopodes” emanating from the distal end of the odontoblast cell bodies in mouse incisor. While the length and thickness (our width) of the “odontopodes” were similar to the dimensions of the plate-like processes we measured, the three dimensional shape was not described. The images and diagrams suggest the presence of a narrowed “neck” at the distal end of the odontoblast, and that the “odontopodes” make a sharp angulation with the cell body, neither of which were evident in our preparations. Additionally, differences in “odontopode” shape related to the position of the odontoblasts along the length of the incisor, or in the molar were not described. Similar to our finding that the plates are aligned in the longitudinal axis of the tooth, it was noted that the “odontopodes” were all oriented toward the tip of the incisor, (Khatibi Shahidi et al 2015).

Our study focused on the structure of odontoblast processes in wild-type mice. Published studies of several genetic mutations demonstrating morphological alterations in dentin suggest that the structure of odontoblast processes may be altered in these mice. In dentin matrix protein 1/Klotho double deficient mice (Rangiani et al., 2012) and mice overexpressing the mutation found in Hutchinson-Gilford Progeria (LMNA, c.1824C > T, p.G608G) in odontoblasts (Choi et al., 2018), dentinal tubules are discontinuous or absent. In mice with mutations in the dentin sialophosphoprotein gene, resin casts of dentin indicate increased thickness and greatly reduced branching of odontoblast processes (Verdelis et al., 2016). Deletion of *Dlx3* in neural crest cells leads to an uneven distribution of thinner and disorganized dentinal tubules (Duverger et al., 2012), and conditional knockout of Raptor/mTORC1 in odontoblasts and mesenchymal cells results in disorganized dentinal tubules (Xie et al., 2019). Disruption of *Smad4* in odontoblasts causes an altered polygonal morphology of root odontoblasts with processes oriented in several directions (Gao et al., 2009). None of these studies, however, examined the three-dimensional structure of the odontoblast processes.

In conclusion, we used serial section electron microscopy to examine the predentin region of the mouse incisor and found unexpected features of odontoblast processes. These processes previously have been depicted as tubules. We confirm this in molars and near the tip of the mouse incisor. However, towards the apex of the incisor, the odontoblast processes are plates that are radially oriented along the apex to tip axis. This shape and orientation may be optimal for the high secretion rates necessary for the continual growth of the incisor. Although tremendous progress has been made in defining the transcription factors and paracrine factors and interactions that govern formation of the tooth, there still remains much to be learned about the cellular structures and processes involved in assembling the tooth.

## Materials and Methods

Wild-type 2 month old male C57BL/6J mice (Jackson Laboratory; Bar Harbor ME), weighing 24-26 grams, were anesthetized by intraperitoneal injection of 0.1 mg ketamine + 0.01 mg xylazine/g body weight. They were fixed by cardiac perfusion with 2.5% glutaraldehyde / 1% paraformaldehyde in 0.1 M sodium cacodylate, pH 7.4, by introducing a needle into the left ventricle of the heart. All protocols were conducted as approved by the University of Connecticut Health Center Animal Care Committee. The mandibles were dissected, and fixed overnight with the same fixation solution. The mandibles were placed in a decalcification solution (257 mM EDTA in PBS, pH 7.4) in a cold room, and checked every 5 days by LX60 digital x-ray machine (Faxitron, Tucson, AZ) to monitor the progress of decalcification, which was complete within 2-3 weeks.

The decalcified incisors and molars were trimmed into 5 segments by using the molars as a reference (Smith and Nanci 1989). Each piece was rinsed several times in buffer, and processed following the ROTO protocol (Tapia et al. 2012). Briefly, the samples were incubated in a reduced osmium solution, treated with a mordant, thiocarbohydrazide, then incubated a second time in unreduced osmium. The samples were then stained overnight in 1% aqueous uranyl acetate at 4°C, then with lead aspartate (Walton 1979) for 30 min. The samples were dehydrated in graded ethanol solutions and infiltrated with graded epoxy resin (Poly/Bed 812, Polysciences, Warrington, PA) / propylene oxide for 14 days before polymerization at 60 °C.

To prepare the samples for sectioning, the block face was shaped using a diamond trimming knife forming a trapezoid shape. The block was mounted in an ultramicrotome (Leica; Buffalo Grove, IL) with the long side of the face oriented vertically. Thin sections (50 – 75 nm in thickness) were cut with a diamond knife (Diatome; Hatfield, PA), and collected with the ATUM tape collector on kapton tape (previously glow discharged in order to reduce wrinkles). The tape with the sections was cut into strips, mounted on a silicon wafer and then carbon coated (Denton 502B; Moorestown, NJ).

The sections were imaged using a field emission scanning electron microscope (Zeiss Sigma FE-SEM, Thornwood, NY or FEI Verios, Hillsboro, OR). Further details on serial sectioning and imaging are described elsewhere (Baena et al. 2019).

The images were aligned using the register virtual stack slices plugin of FIJI Image J. They were segmented using the TrakEM2 module of FIJI Image J and a Cintiq tablet (Wacom; Portland, OR). Reconstructions (.obj format) were viewed using MeshLab (www.meshlab.com) or Photoshop (Adobe; San Jose, CA). To measure distances in the reconstructions, the virtual reality program syGlass (Morgantown, WV) was used with an Oculus Rift headset (Menlo Park, CA).

## Acknowledgements

We thank Valentina Baena for cardiac perfusion, Maya Yankova for sectioning advice, and other members of the Terasaki lab for general support. We also thank Drs. Laurinda Jaffe, Valentina Baena and Rachael Norris for reading the manuscript, and Dr. Charles E. Smith for helpful discussions. NS contributed to the design, data acquisition and interpretation, drafted and critically revised the manuscript. ARH contributed to conception, design and interpretation, drafted and critically revised the manuscript. MT contributed to design and interpretation, drafted and critically revised the manuscript. All authors gave their final approval and agree to be accountable for all aspects of the work. This work was supported by NIH/NIDCR fellowship R90DE022526. The authors declare no potential conflicts of interest with respect to the authorship and/or publication of this article.

## Supplemental files

**Fig2A.mov** Quicktime movie of the serial section images of Figure 2A. In this series, the sections were 50 nm thick. The frames shown in the figure were numbers 100, 111, 122, 134, 145, 156, 168, and 181. The movie speed is 5 frames per second.

**Fig2B.mov** Quicktime movie of the serial section images of Figure 2B. In this series, the sections were 65 nm thick. The frames shown in the figure were numbers 7, 15, 23, 32, 40, 49, 60, and 69. The movie speed is 4 frames per second in order to correspond temporally with panels A and C.

**Fig2C.mov** Quicktime movie of the serial section images of Figure 2C. In this series, the sections were 50 nm thick. The frames shown in the figure were numbers 2, 14, 26, 38, 47, 59, 70, and 81. The movie speed is 5 frames per second.

